# Conserved Ark1-related kinases control TORC2 signaling

**DOI:** 10.1101/870550

**Authors:** Maria Alcaide-Gavilán, Selene Banuelos, Rafael Lucena, Douglas R. Kellogg

## Abstract

In all orders of life, cell cycle progression is dependent upon cell growth, and the extent of growth required for cell cycle progression is proportional to growth rate. Thus, cells growing rapidly in rich nutrients are substantially larger than slow growing cells. In budding yeast, a conserved signaling network surrounding Tor complex 2 (TORC2) controls growth rate and cell size in response to nutrient availability. Here, a search for new components of the TORC2 network identified a pair of redundant kinase paralogs called Ark1 and Prk1. Previous studies found that Ark/Prk play roles in endocytosis. Here, we show that Ark/Prk are embedded in the TORC2 network, where they appear to influence TORC2 signaling independently of their roles in endocytosis. We also show that reduced endocytosis leads to increased cell size, which indicates that cell size homeostasis requires coordinated control of plasma membrane growth and endocytosis. The discovery that Ark/Prk are embedded in the TORC2 network suggests a model in which TORC2-dependent signals control both plasma membrane growth and endocytosis, which would ensure that the rates of each process are matched to each other and to the availability of nutrients so that cells achieve and maintain an appropriate size.

## Introduction

Growth and inheritance are defining features of life. Much is known about inheritance, yet surprisingly little is known about growth. Growth requires myriad biosynthetic processes, all of which must be precisely coordinated with each other and matched to the availability of diverse nutrients. Moreover, the extent of growth must be tightly controlled to ensure that cells maintain an appropriate size.

Not surprisingly, growth is overseen by master regulators that integrate nutrient-dependent signals as well as information from feedback loops to ensure coordination of biosynthetic events. The master regulators of growth in all eukaryotic cells are the Tor kinases, which are assembled into two large multi-protein complexes called TORC1 and TORC2 (Loewith *et al.*, 2002; Wedaman *et al.*, 2003; González and Hall, 2017). TORC1 is inhibited by rapamycin, a cyclic peptide with potent immunosuppressant activity (Heitman *et al.*, 1991). TORC1 has therefore been extensively studied and has well-established roles in control of ribosome biogenesis, translation, nutrient import and autophagy (Wullschleger *et al.*, 2006).

Much less is known about TORC2. In budding yeast, TORC2 directly phosphorylates and activates a pair or redundant kinase paralogs called Ypk1 and Ypk2, which are homologs of vertebrate SGK kinases (Casamayor *et al.*, 1999; Kamada *et al.*, 2005; Niles *et al.*, 2012). Ypk1/2 control production of ceramide, a low abundance lipid that plays roles in signaling (Breslow *et al.*, 2010; Roelants *et al.*, 2011; Muir *et al.*, 2014). Ceramide is also a precursor for production of complex ceramides that serve as components of the plasma membrane. Ypk1/2 control production of ceramide in two ways: they stimulate production of sphingolipids that serve as precursors for ceramide synthesis, and they directly activate ceramide synthase, which builds ceramide from the sphingolipid precursors. The TORC2 signaling network includes a poorly understood negative feedback loop in which ceramide-dependent signals appear to inhibit TORC2 signaling (Roelants *et al.*, 2011; Berchtold *et al.*, 2012; Alcaide-Gavilan *et al.*, 2018; Lucena *et al.*, 2018).

Recent work has shown that the TORC2 network relays nutrient-dependent signals that influence cell size and growth rate (Alcaide-Gavilan *et al.*, 2018; Lucena *et al.*, 2018). In all cells, growth rate is proportional to nutrient availability and cell size is proportional to growth rate (Schaechter *et al.*, 1958; Hirsch and Han, 1969; Johnston *et al.*, 1977; Ferrezuelo *et al.*, 2012; Leitao and Kellogg, 2017). In budding yeast, the TORC2 network is required for these proportional relationships (Lucena *et al.*, 2018; Leitao *et al.*, 2019). TORC2 signaling is high in rich carbon and low in poor carbon. Moreover, reduced activity of Ypk1/2 leads to a large decrease in cell size, and inhibitors of sphingolipid or ceramide synthesis cause dose dependent decreases in cell size and growth rate. Mutant cells that cannot make ceramide show a complete failure in nutrient modulation of growth rate and cell size, as well as a failure in nutrient modulation of TORC2 signaling.

Key questions regarding the TORC2 network remain to be answered. How do nutrients modulate TORC2 signaling? How does ceramide influence growth rate, cell size and TORC2 signaling? What are the downstream targets of the TORC2 network that influence cell growth and size? A better understanding of the TORC2 network will require identification and characterization of additional components of the network.

Here, we searched for additional proteins that play roles in the TORC2 network. A key component of the TORC2 network is Rts1, a conserved regulatory subunit for protein phosphatase 2A. Loss of Rts1 causes a failure in nutrient modulation of cell size, as well as defects in nutrient modulation of TORC2 signaling (Artiles *et al.*, 2009; Lucena *et al.*, 2018; Leitao *et al.*, 2019). Therefore, to identify new components of the TORC2 network we searched for targets of PP2A^Rts1^. In previous work, we compared the phospho-proteomes of wild type and *rts1∆* cells to identify proteins that undergo large changes in phosphorylation in *rts1∆* cells (Zapata *et al.*, 2014). Out of the many thousands of phosphorylation events detected by the mass spectrometry, a subset of 241 sites on 156 proteins were identified as high confidence phosphorylation sites that undergo substantial changes in phosphorylation in *rts1∆* cells. We queried this data set to identify candidate signaling proteins that could work in the TORC2 network. This led to the identification of a kinase called Ark1 as a candidate component of the TORC2 network. Previous studies found that Ark1, along with its redundant paralog Prk1, play roles in controlling late endocytic events (Cope *et al.*, 1999; Zeng *et al.*, 2001; Toshima *et al.*, 2005). Here, we discovered that Ark1 and Prk1 are also required for normal control of TORC2 signaling.

## Results

### The Ark1 and Prk1 kinases are required for nutrient modulation of TORC2 signaling

We first tested whether Ark1 and Prk1 are required for normal signaling in the TORC2 network. TORC2 directly phosphorylates Ypk1 and Ypk2 to stimulate their activity (Kamada *et al.*, 2005). A phosphospecific antibody that recognizes a TORC2 site present on both Ypk1 and Ypk2 therefore provides a readout of TORC2 signaling (Niles *et al.*, 2012; Lucena *et al.*, 2018). We found that loss of Ark1 or Prk1 alone caused no effect, but loss of both caused a reduction in TORC2-dependent signaling to Ypk1/2 (Figure 1A). Loss of Ark/Prk also caused an increase in the amount of Ypk1 protein and in the electrophoretic mobility of Ypk1. Previous work has shown that large changes in Ypk1 electrophoretic mobility are due to the activity Fpk1 and Fpk1, a pair of redundant kinase paralogs that play poorly understood roles in regulation of Ypk1/2 (Roelants *et al.*, 2010).

**Figure 1.**
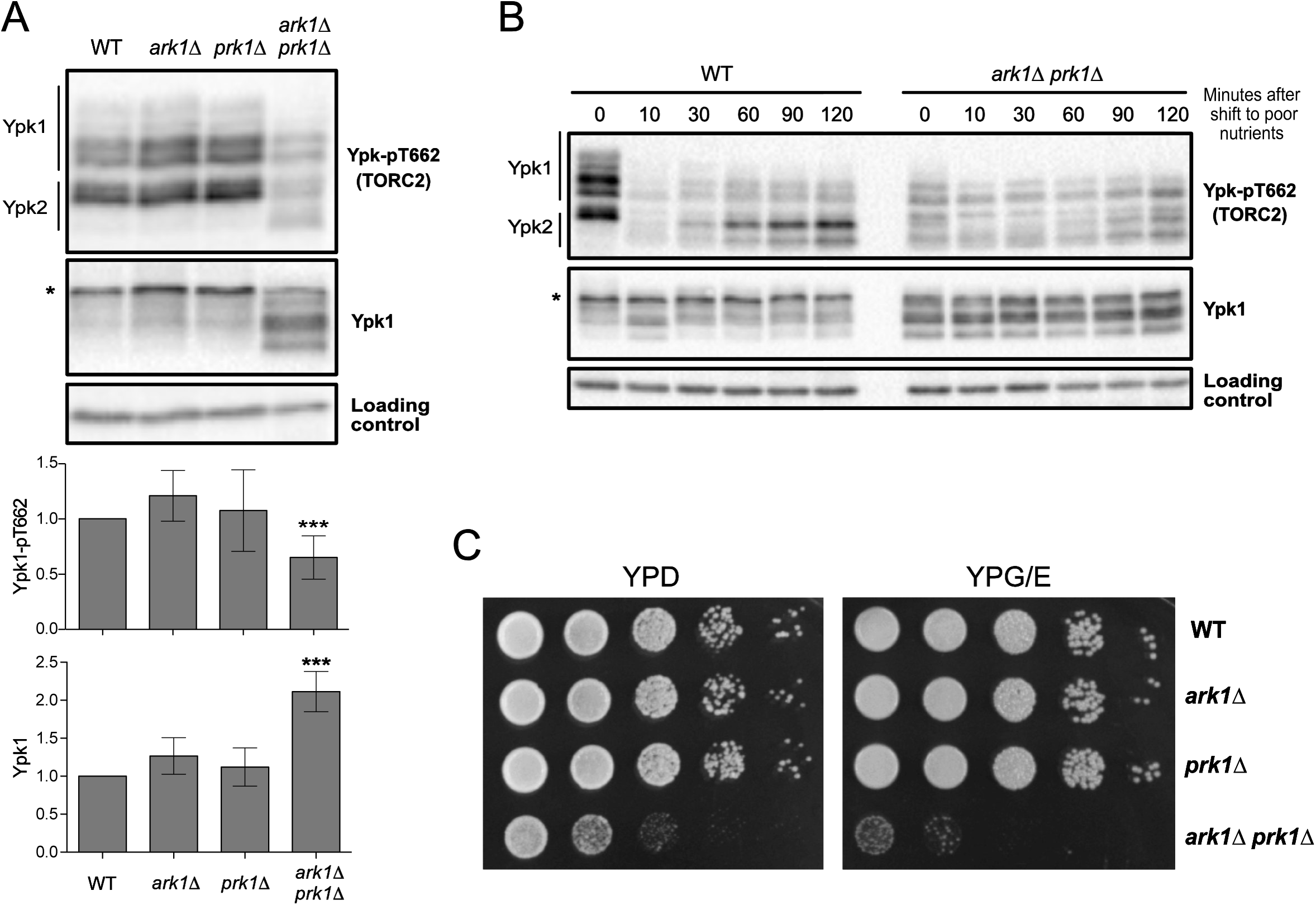
Ark1 and Prk1 are required for nutrient modulation of TORC2 signaling. (A) Cells of the indicated genotypes were grown to log phase in YPD medium at 30°C and levels of Ypk-pT662 and Ypk1 protein were assayed by western blot. An asterisk indicates a background band that overlaps with the Ypk1 signal. Quantification of Ypk-pT662 and total Ypk1 protein are shown. Error bars represent the standard deviation of the mean of three biological replicates. *** indicates a p-value smaller than 0.005 compared to the wildtype. (B) Wildtype and *ark1∆ prk1∆* cells were grown in YPD medium to early log phase and were then rapidly washed into YPG/E medium at 30°C. Cells were collected at the indicated time intervals and western blotting was used to assay levels of Ypk-pT662 and Ypk1 protein. (C) A series of 10-fold dilutions of the indicated strains were grown at 30°C for two days on rich (YPD) or poor (YPG/E) nutrient conditions. A small number of colonies that have suppressor mutations appear when *ark1∆ prk1∆* cells are grown on YPG/E.

We next tested the effects of inhibiting an analog-sensitive allele of *PRK1* in an *ark1∆* background (*prk1-as ark1∆*) (Sekiya-Kawasaki *et al.*, 2003). Inhibition of *prk1-as* caused a reduction in TORC2 signaling within 60 minutes, as well as an increase in Ypk1 protein (Figure S1A).

A protein kinase named Akl1 is a third member of the Ark/Prk kinase family. Analysis of *akl1∆ ark1∆ prk1∆* cells showed that *akl1∆* did not cause additive effects on TORC2 signaling in *ark1∆ prk1∆* cells (Figure S1B).

We next tested whether Ark/Prk are required for modulation of TORC2 signaling in response to changes in carbon source. In wild type cells, a shift from rich carbon (2% glucose) to poor carbon (2% glycerol, 2% ethanol) causes a rapid reduction in TORC2-dependent signaling to Ypk1/2 (Lucena *et al.*, 2018). Over longer times, TORC2 signaling in poor carbon partially recovers, but remains reduced relative to TORC2 signaling in rich carbon. Modulation of TORC2 signaling in response to carbon source failed to occur in *ark1∆ prk1∆* cells (Figure 1B). In addition, *ark1∆ prk1∆* cells fail to proliferate on poor carbon media, consistent with a role for Ark/Prk in the TORC2 signaling network that influences the response to changes in carbon source (Figure 1C).

Together, these observations suggest that Ark/Prk execute functions that are required for normal functioning of the TORC2 network.

### Ark1 and Prk1 respond to nutrient-dependent signals

The TORC2 network responds rapidly to changes in carbon source (Lucena *et al.*, 2018). Therefore, to further test whether Ark1 and Prk1 play roles in TORC2 signaling, we tested whether their phosphorylation state or abundance responds rapidly to changes in carbon source. We first tested whether Ark1 or Prk1 are phosphoproteins. Multiple forms of both Ark1-3XHA and Prk1-6XHA were detected by western blot. Treatment with phosphatase caused the forms to collapse into one band, confirming that Ark1 and Prk1 are phosphorylated (Figure 2A).

**Figure 2.**
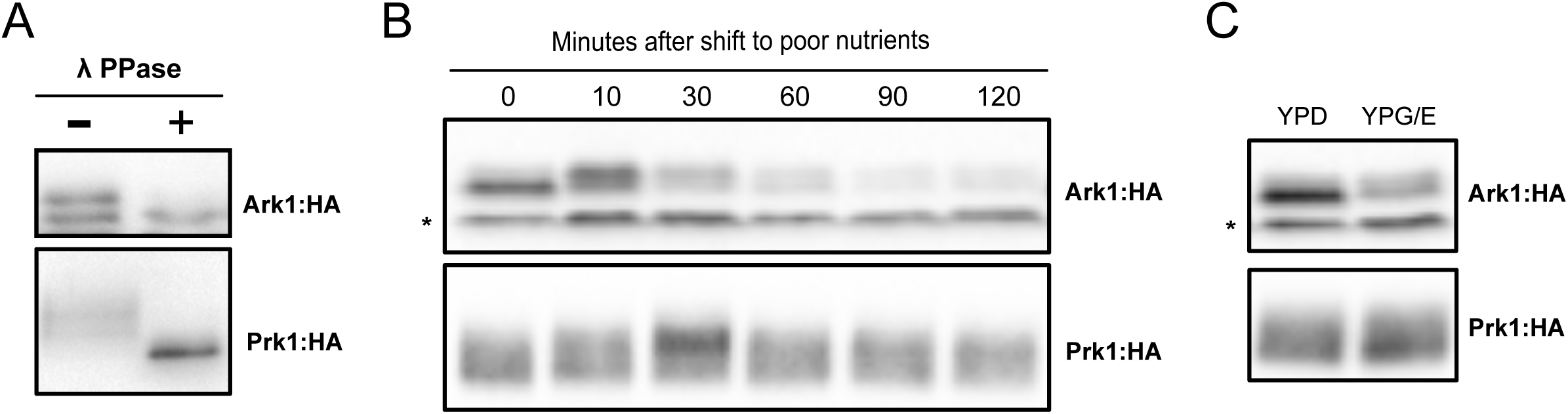
Ark1 and Prk1 respond to nutrient-dependent signals. (A) Extracts from Ark1:3xHA cells and Prk1:6xHA cells were treated with lambda-phosphatase and analyzed by western blot. (B) Ark1:3xHA cells and Prk1:6xHA cells were grown at room temperature in YPD medium to early log phase and were then rapidly washed into YPG/E medium. The behavior of Ark1:3xHA and Prk1:6xHA was assayed by western blot with anti-HA antibody. (C) Ark1:3xHA cells and Prk1:6xHA cells were grown overnight at room temperature in YPD or YPG/E medium and the behavior of Ark1:3xHA and Prk1:6xHA was assayed by western blot. The asterisk indicates a non-specific band.

We next shifted wild type cells from rich to poor carbon and assayed the behavior of both proteins by western blot (Figure 2B). A fraction of Ark1 underwent rapid hyperphosphorylation in response to poor carbon, which was followed by a gradual decrease in protein levels over two hours. Upon prolonged incubation in poor carbon, a fraction of Ark1 persisted in a hyperphosphorylated state and the amount of protein remained reduced (Figure 2C). Prk1 also underwent rapid hyperphosphorylation in response to poor carbon, but showed no change in protein levels. Phosphorylation of Prk1 returned to the starting baseline by the end of the time course. These rapid changes in the phosphorylation state of Ark1 and Prk1 occurred with a timing that was similar to the timing of changes in TORC2 signaling (Figure 1B and (Lucena *et al.*, 2018)).

### Ark1 and Prk1 carry out functions in the TORC2 signaling network

To further investigate the roles of Ark1 and Prk1 in the TORC2 signaling network we tested for functional interactions with components of the network. A simplified model that summarizes proposed functional relationships in the TORC2 network is shown in Figure 3A. Signaling in the network is strongly influenced by feedback loops. Thus, ceramides produced in response to signals from Ypk1/2 relay signals that inhibit TORC2 signaling. The simplified model does not capture all of the data from previous studies. Nevertheless, it provides a framework for interpreting data regarding functional interactions in the network.

**Figure 3.**
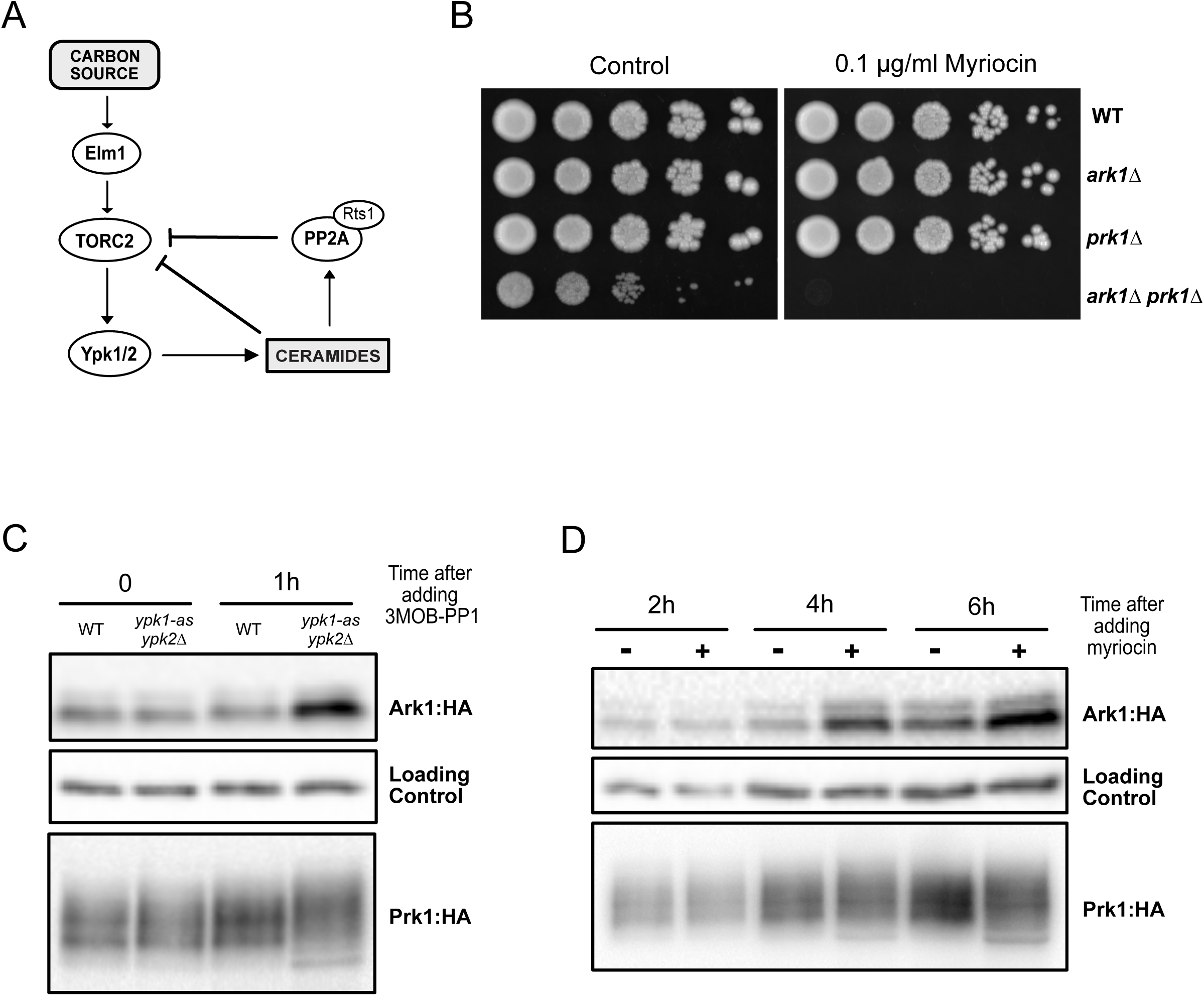
Ark1 and Prk1 carry out functions in the TORC2 signaling network. (A) A simplified model of the TORC2 signaling network. (B) A series of 10-fold dilutions of the indicated strains was grown at 30°C in the presence or absence of 0.1 µg/ml of Myriocin. (C) The indicated strains were grown at room temperature in YPD medium to early log phase and cells were collected before and after one hour after addition of 50 µM 3MOB-PP1. Ark1:3xHA and Prk1:6xHA proteins were analyzed by western blot. (D) Ark1:3xHA and Prk1:6xHA cells were grown at room temperature in YPD medium to early log phase. 1 µM Myriocin or DMSO (control) was added and cells were collected at the indicated times to analyze Ark1:3xHA and Prk1:6xHA proteins by western blot.

We first tested for genetic interactions with Ypk1 and Ypk2. Ypk1 and Ypk2 are partially redundant. Loss of Ypk2 causes little or no phenotype, while loss of Ypk1 causes a large reduction in cell size and a reduced rate of proliferation (Lucena *et al.*, 2018). We were unable to recover *ypk1∆ ark1∆ prk1∆* triple mutants. We also found that *ark1∆ prk1∆* cells are hypersensitive to myriocin, an inhibitor of the enzyme serine palmitoyltransferase, which catalyzes the first step in production of sphingolipids that are required for normal signaling in the TORC2 network (Figure 3B). These interactions are consistent with a role for Ark1 and Prk1 in promoting signaling in the TORC2 network.

We next tested whether inhibition of Ypk1/2 causes effects on Ark1 or Prk1. To do this, we used an analogue sensitive allele of Ypk1 in an *ypk2∆* background (*ypk1-as ypk2Δ*) (Sun *et al.*, 2012). Inhibition of *ypk1-as* caused an increase in Ark1 protein levels and a change in Prk1 phosphorylation within one hour (Figure 3C).

Ypk1/2 promotes synthesis of ceramide from sphingolipid precursors, and ceramide is required for feedback signals that repress TORC2 signaling (Roelants *et al.*, 2011; Muir *et al.*, 2014; Lucena *et al.*, 2018). Therefore, we next tested how inhibiting production of sphingolipids and ceramide influences the Ark1 and Prk1 proteins. We found that addition of myriocin caused the same effects as inhibition of *ypk1-as* (Figure 3D). Previous work found that myriocin and inhibition of Ypk1/2 both cause increased TORC2 activity, whereas poor carbon causes reduced TORC2 activity (Lucena *et al.*, 2018). Thus, changes in the abundance and phosphorylation of the Ark1 protein are correlated with TORC2 activity.

We also tested for functional relationships with PP2A^Rts1^. Loss of function of Rts1 causes increased signaling in the TORC2 network (Lucena *et al.*, 2018). Here, we found that increased TORC2 signaling in *rts1∆* cells is dependent upon Ark1 and Prk1 (Figure 4A). We further found that *rts1∆* enhanced the slow proliferation phenotype caused by *ark1∆ prk1∆* (Figure 4B).

**Figure 4.**
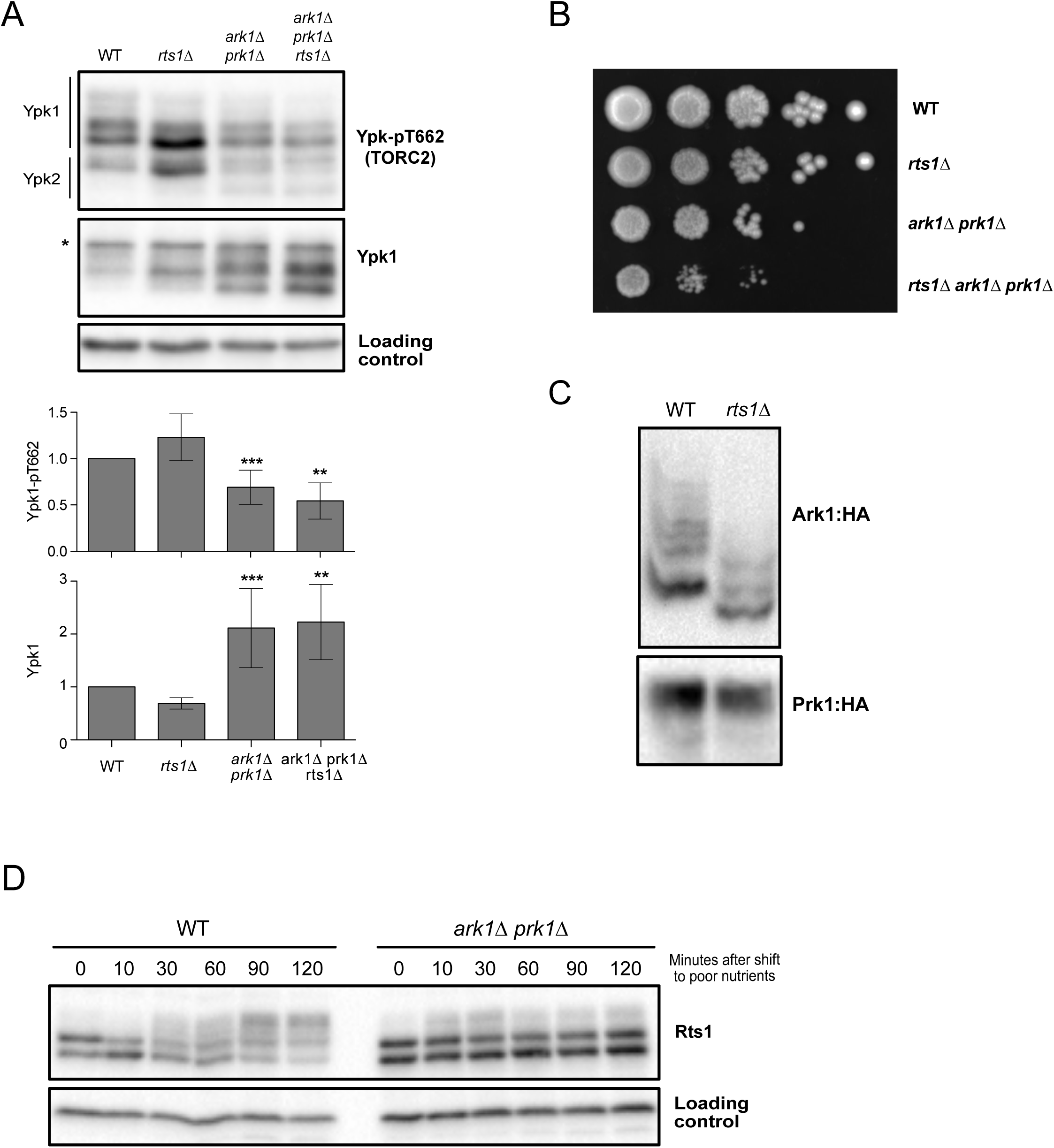
Ark1 and Prk1 show genetic interactions with Rts1. (A) Cells of the indicated genotypes were grown at room temperature to log phase in YPD medium. Ypk-pT662 and total Ypk1 were assayed by western blot. An asterix indicates a background band that overlaps with the Ypk1 signal. Quantifications of Ypk-T662 phosphorylation and total Ypk1 protein are shown. Error bars represent the SD of the mean of three biological replicates. *** indicates a p-value smaller than 0.005; ** indicates a p-value between 0.01 and 0.005 when the indicated strain is compared to the wildtype. (B) A series of 10-fold dilutions of the indicated strains were grown at 25°C on YPD medium. (C) Wildtype and *rts1∆* cells were grown to log phase in YPD medium and phosphorylation of Ark1:3xHA and Prk1:6xHA was assayed by PhosTag western blot. (D) Wild-type and *ark1∆ prk1∆* kinase mutants were grown at 30°C in YPD medium and were then shifted to YPG/E medium. Rts1 was detected by Western blotting.

Since our previous mass spectrometry analysis suggested that Ark1 could be a target of PP2A^Rts1^, we also analyzed the phosphorylation state of Ark1 and Prk1 in *rts1∆* cells. No change in phosphorylation was observed by normal western blot in log phase cells or in nutrient shifted cells (Figure S2A and B). However, analysis of phosphorylation using Phos-tag gels in rapid growing cells and nutrient shifted cells showed a loss of Ark1 phosphorylation in *rts1∆* cells, consistent with the mass spectrometry data (Figure 4C and Figure S2C). No changes in phosphorylation of Prk1 protein were observed.

In previous work, we found that Rts1 undergoes TORC2-dependent hyperphosphorylation in response to a shift to poor carbon (Alcaide-Gavilan *et al.*, 2018). Here, we found that hyperphosphorylation of Rts1 in response to poor carbon was strongly reduced in *ark1∆ prk1∆* cells (Figure 4D). The effects of *ark1∆ prk1∆* on Rts1 hyperphosphorylation were similar to the effects caused by inactivation of Ypk1/2 or other components of the TORC2 network.

Elm1, a budding yeast homolog of vertebrate LKB1 kinase, is required for normal levels of TORC2 signaling and for nutrient modulation of TORC2 signaling (Alcaide-Gavilan *et al.*, 2018). We tested whether Elm1 is required for nutrient-dependent signaling to Ark1 and Prk1. We found that phosphorylation of Ark1 and Prk1 in response to poor carbon is not dependent upon Elm1 (Figure S2D and E). We also found that *elm1∆* is synthetically lethal with *ark1∆ prk1∆*, which suggests that Elm1 could promote TORC2 signaling via a pathway that works in parallel with Ark/Prk.

Together, the data show that Ark/Prk play roles in the TORC2 signaling network.

### Cell size defects in *ark1∆ prk1∆* cells are likely caused by defects in endocytosis

Ark/Prk kinases are known to regulate endocytic vesicle uncoating via phosphorylation of multiple coat proteins, including Pan1, Ent1 and Sla1 (Watson *et al.*, 2001; Zeng *et al.*, 2001). Previous work has shown that endocytic proteins Ent1 and Pan1 are TORC2/Ypk1 effectors (Rispal *et al.*, 2015). Since ark*1∆ prk1∆* cells are abnormally large, we reasoned that their increased size could be due to defects in TORC2 signaling, defects in endocytosis, or both. In previous work, we showed that defects in TORC2 signaling causes defects in nutrient modulation of cell size, as well as defects in cell size modulation in response to sub-lethal doses of myriocin (Lucena *et al.*, 2018). However, the discovery that *ark1∆ prk1∆* cells fail to proliferate in poor carbon sources or in the presence of low doses of myriocin made it impossible to test whether they show normal cell size responses to these conditions.

To determine whether the large size of *ark1∆ prk1∆* cells could be due to defects in endocytosis, we tested whether reduced endocytosis causes increased cell size. Ent1 and Ent2 are redundant paralogs that drive early steps in endocytosis (Wendland *et al.*, 1999). To inactivate Ent1/2, we generated an auxin-inducible degron version of *ENT2* in an *ent1∆* background (*ent2-AID ent1∆*). The *ent2-AID ent1∆* cells were viable in the absence of auxin and failed to proliferate in the presence of 15 µM auxin (Figure S3A). The *ent2-AID* protein disappeared within 10 minutes of adding auxin (Figure S3B).

In the absence of auxin, *ent2-AID ent1∆* cells were abnormally large, similar to the increase caused by *ark1∆ prk1∆* (Figure 5). Sub-lethal doses of auxin caused *ent2-AID ent1∆* cells to become even larger, suggesting that reduced endocytosis strongly influences cell size. Thus, the large size of *ark1∆ prk1∆* cells may be due largely to defects in endocytosis. The increase in cell size caused by defects in endocytosis suggests that control of cell size requires precise coordination of exocytosis and endocytosis.

**Figure 5.**
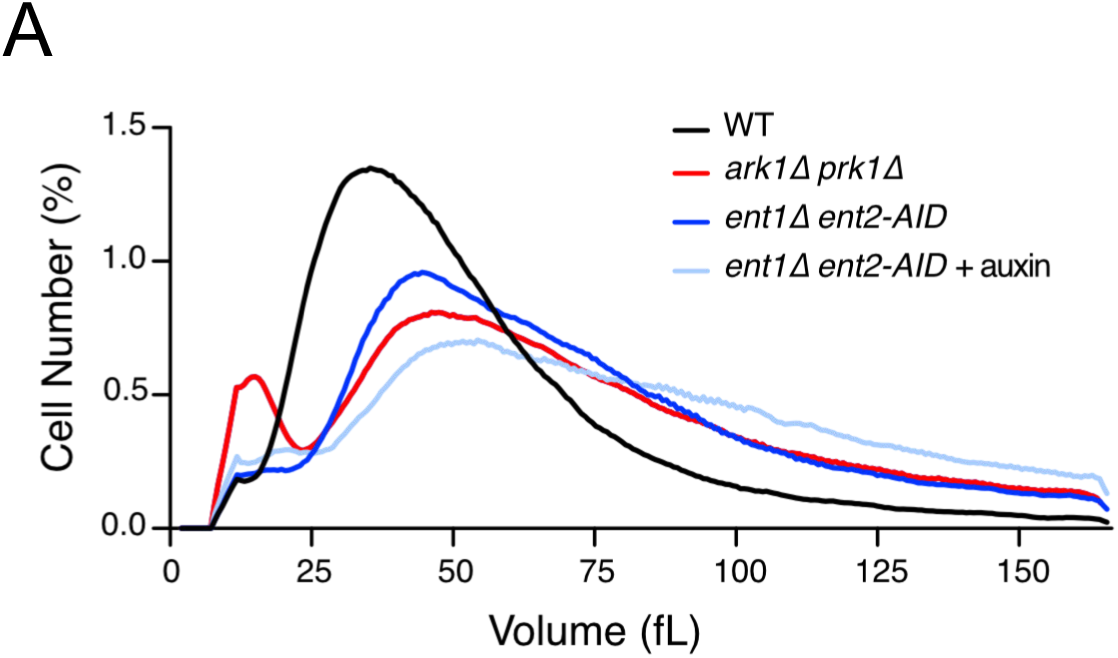
Cell size defects in *ark1∆ prk1∆* cells are likely caused by defects in endocytosis. (A) Cells of the indicated genotypes were grown to log phase at 25°C and cell-size distributions were determined using a Coulter counter. *ent2-AID ent1∆* cells were grown in the absence or presence of 5 µM auxin. Each plot is the average of 3 biological replicates. For each biological replicate, 3 technical replicates were analyzed and averaged.

### The effects of Ark/Prk on TORC2 signaling are unlikely to be a consequence of reduced endocytosis

Since Ark/Prk are required for normal endocytosis, we tested whether defects in TORC2 signaling caused by *ark1∆ prk1∆* could be a consequence of defects in endocytosis. To do this, we examined how mutants in several endocytic proteins that are targets of Prk1 influence TORC2 signaling. We first tested the effects of loss of function of Ede1, which is required for efficient execution or early endocytic steps. Previous work found that *ede1∆* causes an 35% reduction in endocytosis (Gagny *et al.*, 2000). We found that *ede1∆* had no effect on the level of TORC2 signaling in unsynchronized cells growing in rich carbon, but caused an increase in the amount of Ypk1 protein (Figure 6A). We also found that *ede1∆* had no effect on the response of the TORC2 network to a shift from rich to poor carbon (Figure 6B). We next tested essential components of the endocytic machinery. Inactivation of Ent1/2 in rapidly growing cells had no effect on the level of TORC2-dependent phosphorylation of Ypk1/2, but did cause an increase in levels of the Ypk1 protein (Figure 6C). Growing cells overnight in the presence of a semi-lethal dose of auxin that causes an increase in cell size (Figure 5) also had no effect on TORC2 signaling (**Fig S4A**). Moreover, inactivation of Ent1/2 30 minutes before a shift to poor carbon had no effect on modulation of TORC2 signaling in response to a shift to poor carbon (Figure 6D). Finally, we observed that *ede1∆* and *ent2-AID ent1∆* mutants are more resistant to myriocin, contrary to what we observed for *ark1∆ prk1∆* cells (**Figures S4B and 3B**).

**Figure 6.**
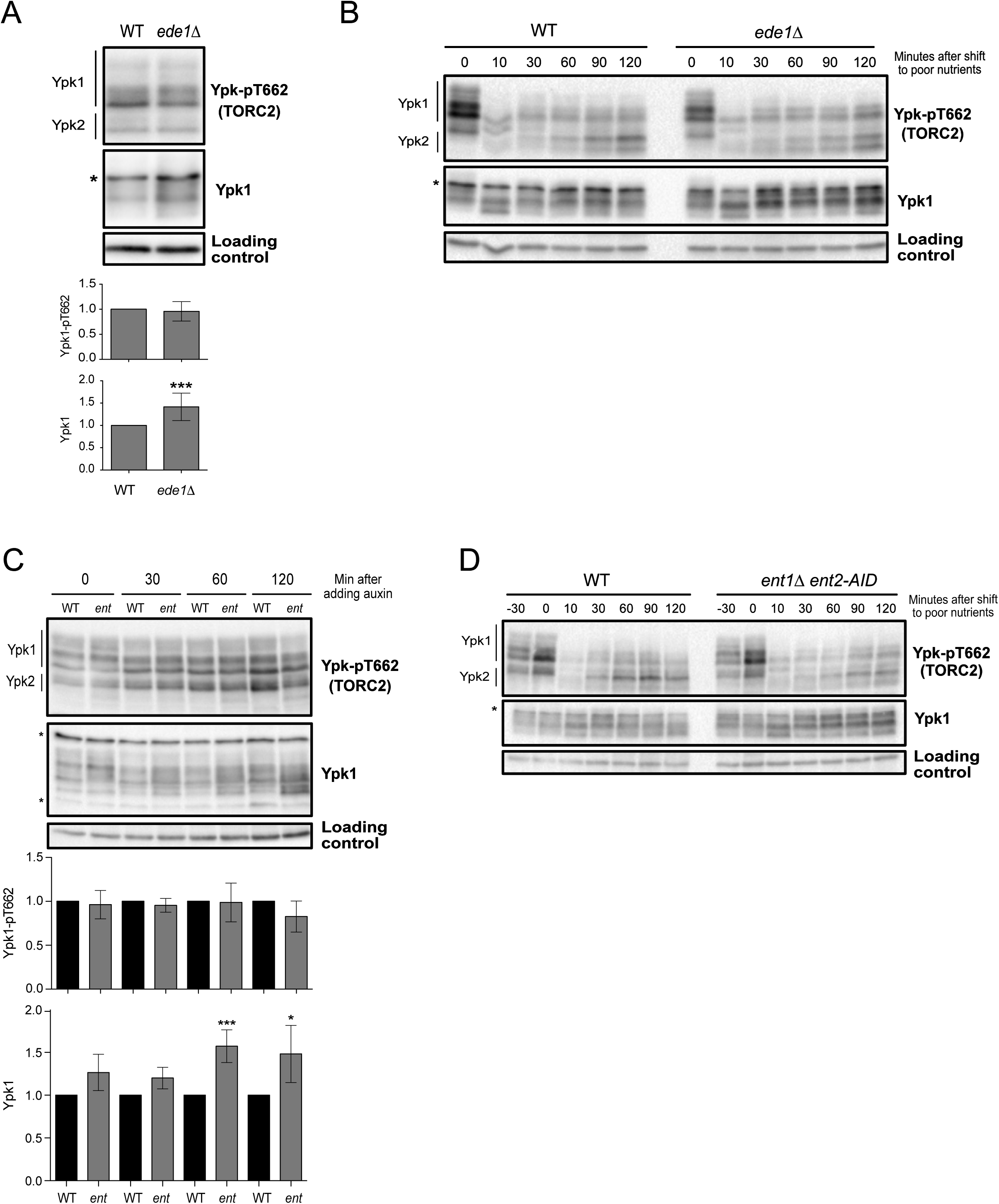
Early and intermediate endocytic mutants have no effects on TORC2 signaling. (A) Wild type and *ede1∆* cells were grown at 30°C to log phase in YPD medium and Ypk-pT662 and Ypk1 protein were assayed by western blot. Quantification of Ypk-T662 phosphorylation and total Ypk1 protein are shown. (B) Wild type and *ede1∆* cells were grown in YPD medium to early log phase and were then rapidly washed into YPG/E medium at 30°C. Cells were collected at the indicated time intervals and levels of Ypk-pT662 and Ypk1 protein were assayed by western blot. An asterisk indicates a background band that overlaps with the Ypk1 signal. (C) Cells of the indicated genotypes were grown at 30°C to log phase in YPD medium. 1 mM Auxin was added and cells were collected and analyzed by western blot at the indicated time points. Quantification of Ypk-T662 phosphorylation and total Ypk1 protein are shown. An asterisk indicates a background band that overlaps with the Ypk1 signal. (D) Cells of the indicated genotypes were grown in YPD medium to early log phase. 1 mM Auxin was added and after 30 minutes cells were washed into YPG/E medium at 30°C containing 1 mM Auxin. Cells were collected at the indicated time intervals and Ypk-pT662 and Ypk1 were assayed by western blot. An asterisk indicates a background band that overlaps with the Ypk1 signal. Error bars represent the standard deviation of the mean of three biological replicates. *** indicates a p-value smaller than 0.005; ** indicates a p-value between 0.01 and 0.005 and indicates a p-value between 0.05 and 0.01 when the indicated strain is compared to the wildtype.

A key target of Ark/Prk in the endocytic pathway is Pan1, an epsin-like protein that promotes polymerization of actin to help drive formation and internalization of endocytic vesicles. There are 19 potential Ark/Prk phosphorylation sites in Pan1 and it has been demonstrated that Ark/Prk directly phosphorylate and inhibit Pan1 (Toshima *et al.*, 2005; 2016). Since Pan1 is an essential player in the last steps of endocytosis, we tested whether loss of Pan1 could have an effect in TORC2 signaling. We used an auxin-inducible degron version of Pan1 to analyze how inactivation of Pan1 influences TORC2 signaling (Bradford *et al.*, 2015). Inactivation of pan1-AID did not have a significant effect on TORC2 signaling in cells growing in rich carbon or in cells shifted from rich to poor carbon (Figures 7A and B). Inactivation of pan1-AID caused increased levels of Ypk1 protein (Figure 7A).

**Figure 7.**
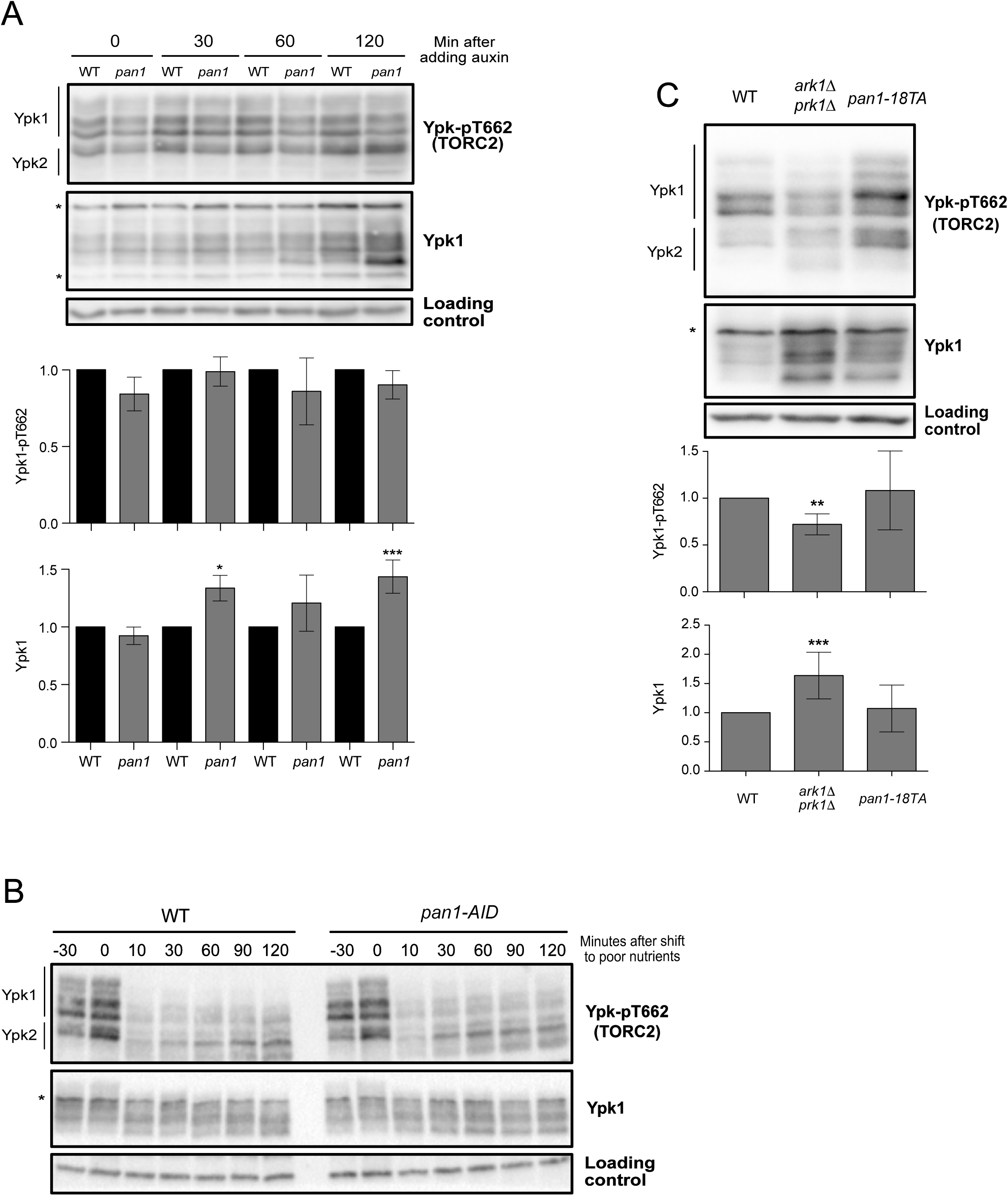
Pan1 mutants do not show significant defects in TORC2 signaling. (A) Wild type and *pan1-AID* cells were grown at 30°C to log phase in YPD medium. 1 mM Auxin was added and cells were collected at the indicated intervals and Ypk-pT662 and Ypk1 protein levels were analyzed by western blot. Quantification of Ypk-T662 phosphorylation and total Ypk1 protein are shown. (B) Wild type and *pan1-AID* cells were grown in YPD medium to early log phase. 1 mM Auxin was added and after 30 minutes cells were washed into YPG/E medium at 30°C containing 1 mM Auxin. Cells were collected at the indicated intervals and Ypk-pT662 and Ypk1 protein levels were assayed by western blot. (C) Cells of the indicated genotypes were grown in YPD medium to early log phase at 30°C and Ypk-pT662 and Ypk1 protein were assayed by western blot. An asterisk indicates a background band in the Ypk1 blot that overlaps with the Ypk1 signal. Error bars represent the standard deviation of the mean of three biological replicates. *** indicates a p-value smaller than 0.005 and ** indicates a p-value between 0.01 and 0.005 when the indicated strain is compared to the wildtype.

We next considered the possibility that the effects of *ark1∆ prk1∆* on TORC2 signaling are due to hyperactivity of Pan1. Previous work found that a phospho-site mutant of Pan1 (*pan1-18TA*) that cannot be phosphorylated by Ark/Prk phenocopies the effects of *ark1∆ prk1∆* on actin organization and endocytosis (Toshima *et al.*, 2016). We found that the *pan1-18TA* mutant had no effect on TORC2 signaling or total Ypk1 protein. (Figure 7C).

Since *ark1∆ prk1∆* do not proliferate on poor carbon (Figure 1C) we tested whether endocytic mutants influence growth on poor carbon. However, we found that *ede1∆* and *pan1-18TA* had no effect on proliferation on poor carbon (**Figures S4C, E**). Growth of *ent2-AID ent1∆* or *pan1-AID* in the presence of sublethal doses of auxin also had no effect on proliferation in poor carbon (Figure S4B and S4D).

Together, the data suggest that Ark/Prk influence the level of TORC2 signaling independently of their role in endocytosis. Furthermore, defects in endocytosis, caused by loss of Ark/Prk or endocytic proteins, cause increased levels of Ypk1 protein.

## Discussion

Previous studies suggested that Ark1 and Prk1 function at a late stage of endocytosis to inhibit actin polymerization and promote disassembly of endocytic coat proteins (Cope *et al.*, 1999; Watson *et al.*, 2001; Zeng *et al.*, 2001; Henry *et al.*, 2003; Toshima *et al.*, 2005; 2016). Here, we show that Ark/Prk are also required for normal control of TORC2 signaling. Loss of Ark/Prk causes reduced TORC2 signaling, as well as a failure in modulation of TORC2 signaling in response to carbon source. Ark/Prk also show strong genetic interactions with multiple components of the TORC2 signaling network. Thus, *ark1∆ prk1∆* are synthetically lethal with loss of *YPK1* or *ELM1*, both of which are required for normal levels of TORC2 signaling. In addition, *ark1∆ prk1∆* cells are exquisitely hypersensitive to myriocin, which inhibits production of sphingolipids that relay signals in the TORC2 network. Loss of Rts1, an important regulator of the TORC2 network causes a loss of Ark1 phosphorylation. In addition, *rts1∆* causes increased signaling in the TORC2 network that is dependent upon Ark/Prk, and hyperphosphorylation of Rts1 in response to a shift from rich to poor carbon is dependent upon Ark/Prk.

Additional observations show that the Ark1 and Prk1 proteins respond to changes in carbon source, consistent with a role in TORC2 signaling. Ark1 shows rapid and sustained changes in both abundance and phosphorylation in response to a switch from rich to poor carbon. Prk1 also shows changes in phosphorylation. Furthermore, *ark1∆ prk1∆* cells are inviable when grown on a poor carbon source, which indicates that Ark1 and Prk1 relay signals that are essential for survival in poor carbon. A model that is consistent with the functional interactions that we observed between the TORC2 network and Ark/Prk is shown in Figure 8.

**Figure 8.**
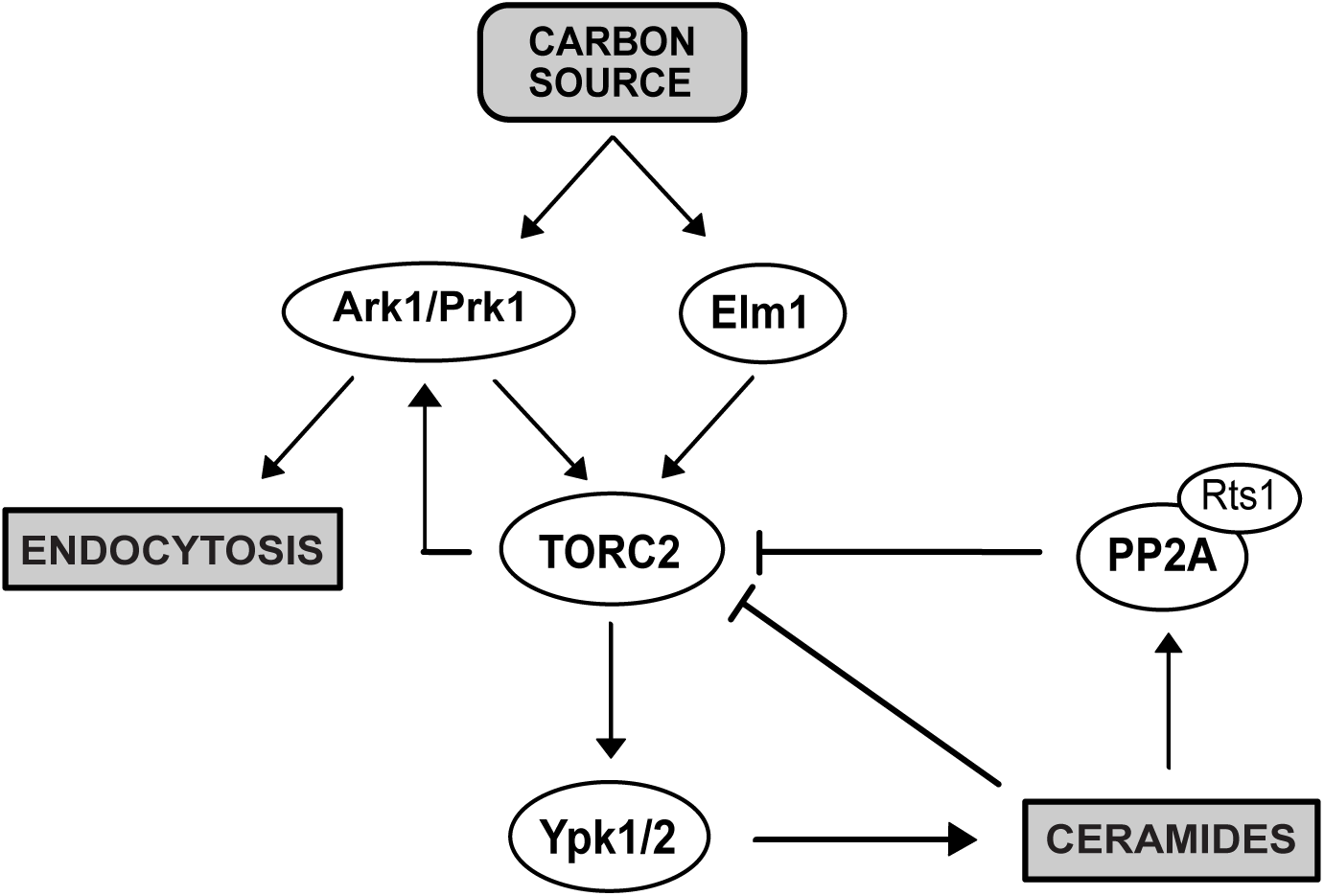
A model summarizing hypothesized functional interactions between Ark-family kinases and the TORC2 signaling network.

A number of observations suggest that Ark/Prk do not influence TORC2 signaling solely via their roles in endocytosis. For example, loss of Ark/Prk causes decreased TORC2 signaling and increased Ypk1 levels, whereas loss of endocytosis only causes increased Ypk1 levels. In addition, a Pan1 phosphorylation site mutant that lacks all of the sites targeted by Ark/Prk phenocopies the effects of *ark1∆ prk1∆* on endocytosis yet causes no effects on TORC2 signaling. Finally, the inviability of *ark1∆ prk1∆* cells on poor carbon is difficult to explain if one assumes that Ark/Prk function solely in endocytosis, since no previous studies have found that efficient endocytosis is required for survival on poor carbon. Together, these observations suggest that Ark/Prk relay signals that influence TORC2 signaling independently of their roles in endocytosis.

An interesting possibility is that Ark/Prk are embedded in the TORC2 network, where they influence TORC2 signaling, while also relaying TORC2-dependent signals that influence endocytosis. Previous studies found that TORC2-dependent signals are required for endocytosis. For example, inhibition of TORC2 activity causes a rapid block to endocytosis (Rispal *et al.*, 2015). Similarly, inhibiting synthesis of sphingolipids that are required for TORC2-dependent signals causes a rapid block to endocytosis (Zanolari *et al.*, 2000).

Our discovery that reduced endocytosis leads to increased cell size suggests that plasma membrane growth and endocytosis must be precisely coordinated to ensure that the two processes are appropriately balanced to achieve and maintain an appropriate cell size. Furthermore, when cells are shifted from rich to poor carbon, the rates of biosynthetic processes that drive plasma membrane growth decrease, so it is likely that the rate of endocytosis must also be decreased to ensure that cells reach an appropriate size. Having common TORC2-dependent signals control both membrane growth and endocytosis could ensure that the rates of each process are appropriately matched to each other and to the availability of nutrients so that cells maintain an appropriate cell size.

## Materials and Methods

### Yeast strains and media

All strains are in the W303 background (*leu2-3,112 ura3-1 can1-100 ade2-1 his3-11,15 trp1-1 GAL+ ssd1-d2*), with the exception of strains DDY903, DDY2544 and JJTY0888 used in Figures 7C and S4E, which is in the S288C background (*his3-∆200, ura3-52, lys2-801, leu2-3, 112*,). Table 1 shows additional genetic features. One-step PCR-based gene replacement was used for making deletions and adding epitope tags at the endogenous locus (Longtine, 1998; Janke et al., 2004). Cells were grown in YP medium (1% yeast extract, 2% peptone, 40 mg/L adenine) supplemented with 2% dextrose (YPD) or 2% glycerol/ethanol (YPG/E). For nutrient shifts, cells were grown in YPD medium overnight to log phase. Cells were then washed 3 times with YPG/E medium and resuspended in YPG/E medium.

Myriocin (Sigma) was dissolved in 100% methanol to make a 500 µg/ml stock solution. The same volume of methanol was added in control experiments. We observed significant differences in the effective concentration of myriocin in different batches from the same supplier.

### Production of polyclonal Ypk1 antibody

An antibody that recognizes Ypk1 was generated by immunizing rabbits with a fusion protein expressed in bacteria. Briefly, a PCR product that includes the full-length open reading frame for Ypk1 was cloned into the entry vector pHis::parallel. The resulting plasmid expresses Ypk1 fused at its N-terminus to 6XHis-TEV. The 6XHis-TEV-Ypk1 fusion was expressed in Rosetta cells and purified using standard procedures, yielding 80 mg of protein from 3 L of bacterial culture. A milligram of the purified protein was used to immunize a rabbit. The 6XHis-TEV-Ypk1 fusion protein was coupled to Affigel 10 (Bio-Rad) to create an affinity column for purification of the antibody.

### Western blotting

For western blots using cells growing in early log phase, cultures were grown overnight at the specified temperature to an OD_600_ of less than 0.8. After adjusting optical densities to normalize protein loading, 1.6-ml samples were collected and centrifuged at 13,000 rpm for 30 s. The supernatant was removed and 150 µl of glass beads were added before freezing in liquid nitrogen.

To analyze cells shifted from rich to poor nutrients, cultures were grown in YPD overnight at 25°C to an OD_600_ of less than 0.8. After adjusting optical densities to normalize protein loading, cells were washed three times with the same volume of YPG/E medium and then incubated at 30°C in YPG/E for the time course. 1.6-ml samples were collected at each time point.

Cells were lysed into 140 μl of sample buffer (65 mM Tris-HCl, pH 6.8, 3% SDS, 10% glycerol, 50 mM NaF, 100 mM β -glycerophosphate, 5% 2-mercaptoethanol, and bromophenol blue). PMSF was added to the sample buffer to 2 mM immediately before use. Cells were lysed in a Mini-bead-beater 16 (BioSpec) at top speed for 2 min. The samples were removed and centrifuged for 15 s at 14,000 rpm in a microfuge and placed in boiling water for 5 min. After boiling, the samples were centrifuged for 5 min at 15,000 rpm and loaded on an SDS polyacrylamide gel.

Samples were analyzed by western blot as previously described (Harvey *et al.*, 2011). SDS-PAGE gels were run at a constant current of 20 mA and electrophoresis was performed on gels containing 10% polyacrylamide and 0.13% bis-acrylamide. Proteins were transferred to nitrocellulose using a Trans-Blot Turbo system (Bio-Rad Laboratories). Blots were probed with primary antibody overnight at 4°C. Proteins tagged with the HA epitope were detected with the 12CA5 anti-HA monoclonal antibody (gift of David Toczyski, University of California San Francisco) or an anti-HA polyclonal antibody developed in our lab. Rabbit anti-phospho-T662 antibody (gift of Ted Powers, University of California, Davis) was used to detect TORC2-dependent phosphorylation of YPK1/2 at a dilution of 1:20,000 in TBST (10 mM Tris-Cl, pH 7.5, 100 mM NaCl, and 0.1% Tween 20) containing 3% milk. Total Ypk1 was detected using anti-Ypk1 antibody (Santa Cruz Biotechnology, catalog number sc-12051, not produced anymore) at a dilution of 1:2000 or the Ypk1 polyclonal antibody previously described. V5 Tag monoclonal antibody (E10/V4RR, Thermo-Fisher MA5-15253) was used to detect *ent2-AID* at 1:2500 in PBS containing 3% milk.

All blots were probed with an HRP-conjugated donkey anti-rabbit secondary antibody (GE Healthcare, catalog number NA934V) or HRP-conjugated donkey anti–mouse antibody (GE Healthcare, catalog number NXA931) or HRP-conjugated donkey anti-goat (Santa Cruz Biotechnology, catalog number sc-2020) for 45–90 min at room temperature. Secondary antibodies were detected via chemiluminescence with Advansta ECL reagents.

Densitometric quantification of western blot signals was performed using ImageJ(Schneider *et al.*, 2012). Quantification of Ypk-pT662 phosphorylation was calculated as the ratio of the phospho-specific signal over the total Ypk1 protein signal, with wild-type signal normalized to a value 1. At least three biological replicates were analyzed and averaged to obtain quantitative information.

### Lambda phosphatase treatment

Lambda phosphatase treatment of Ark1 and Prk1 in cell extracts was carried out as previously described (Lucena *et al.*, 2017).

## Abbreviations used

TORC2: Target of Rapamycin 2
HA: Human Influenza Hemagglutinin
AID: Auxin Inducible Degron

## Data Availability

Strains and plasmids are available upon request. The authors affirm that all data necessary for confirming the conclusions of the article are present within the article, figures, and tables.

## Acknowledgments

We thank members of the laboratory for advice and support. We also thank Ted Powers (UC Davis) for the Ypk-pT662 phosphospecific antibody and David Toczyski (UCSF) for 12CA5 antibody. We are grateful to Yidi Sun and Beverly Wendland for strains and Carrie Partch (UCSC) for pHis::parallel plasmid. This work was supported by National Institutes of Health grant GM053959.

---

**Supplementary Figure 1.**
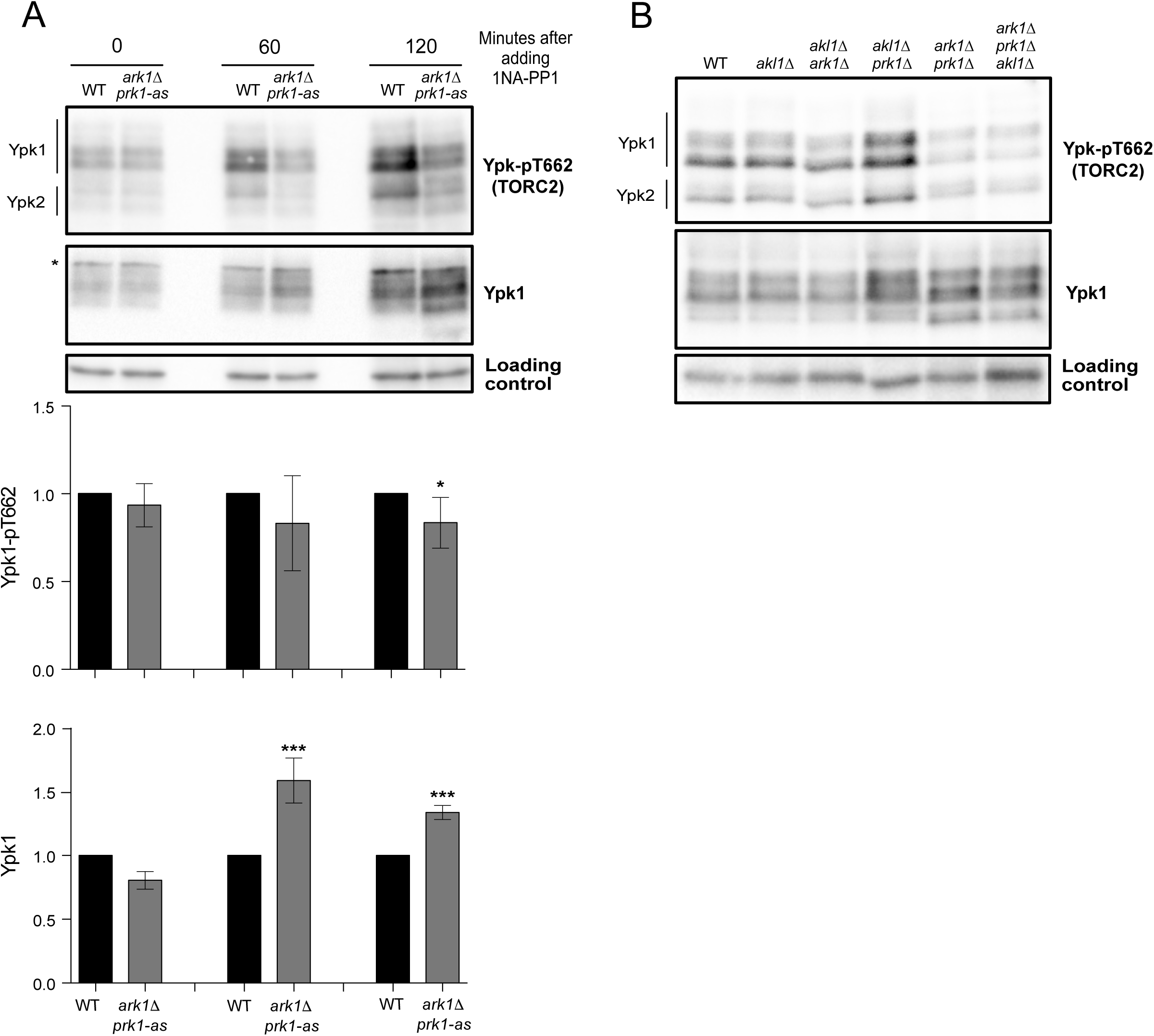
(A) Wildtype and *ark1∆ prk1-*as cells were grown to log phase in YPD medium at 30°C and 80 µM 1NA-PP1 was added to both strains. Cells were collected at the indicated time intervals and Ypk-pT662 and Ypk1 protein were assayed by western blot. Error bars represent the standard deviation of the mean of three biological replicates. *** indicates a p-value smaller than 0.005; ** indicates a p-value between 0.01 and 0.005 and * indicates a p-value between 0.05 and 0.01 when the indicated strain is compared to the wildtype (B) Cells of the indicated genotypes were grown to log phase in YPD medium at 30°C and Ypk-pT662 and Ypk1 protein were assayed by western blot.

**Supplementary Figure 2.**
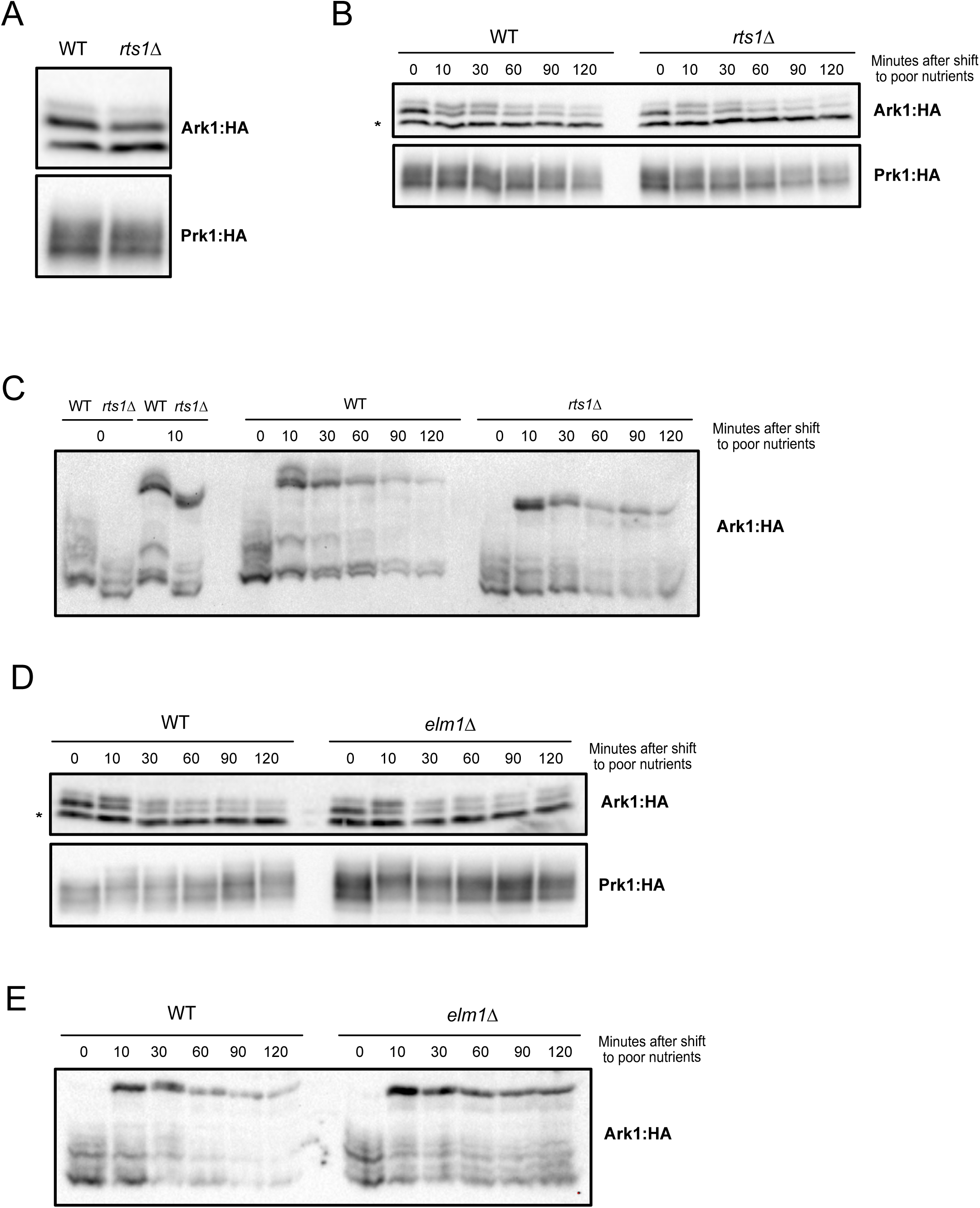
(A) Wildtype and *rts1∆* cells were grown to log phase in YPD medium at 30°C and phosphorylation of Ark1:3xHA and Prk1:6XHA was analyzed by western blot. (B) Wildtype and *rts1∆* mutant cells grown in YPD medium to early log phase and were then rapidly washed into YPG/E medium at 30°C. Cells were collected at the indicated time intervals and phosphorylation of Ark1:3xHA and Prk1:6XHA was analyzed by western blot. An asterisk indicates a background band. (C) Same as (B), but phosphorylation was assayed by PhosTag western blot. On the left, wildtype and *rts1∆* cells were directly compared 10 minutes after a shift to YPG/E. (D) Wildtype and *elm1∆* cells were grown in YPD medium to early log phase and were then rapidly washed into YPG/E medium at 30°C. Cells were collected at the indicated time intervals and phosphorylation of Ark1:3xHA and Prk1:6XHA was analyzed by western blot. An asterisk indicates a background band. (E) Same as D, but phosphorylation was assayed by PhosTag western blot.

**Supplementary Figure 3.**
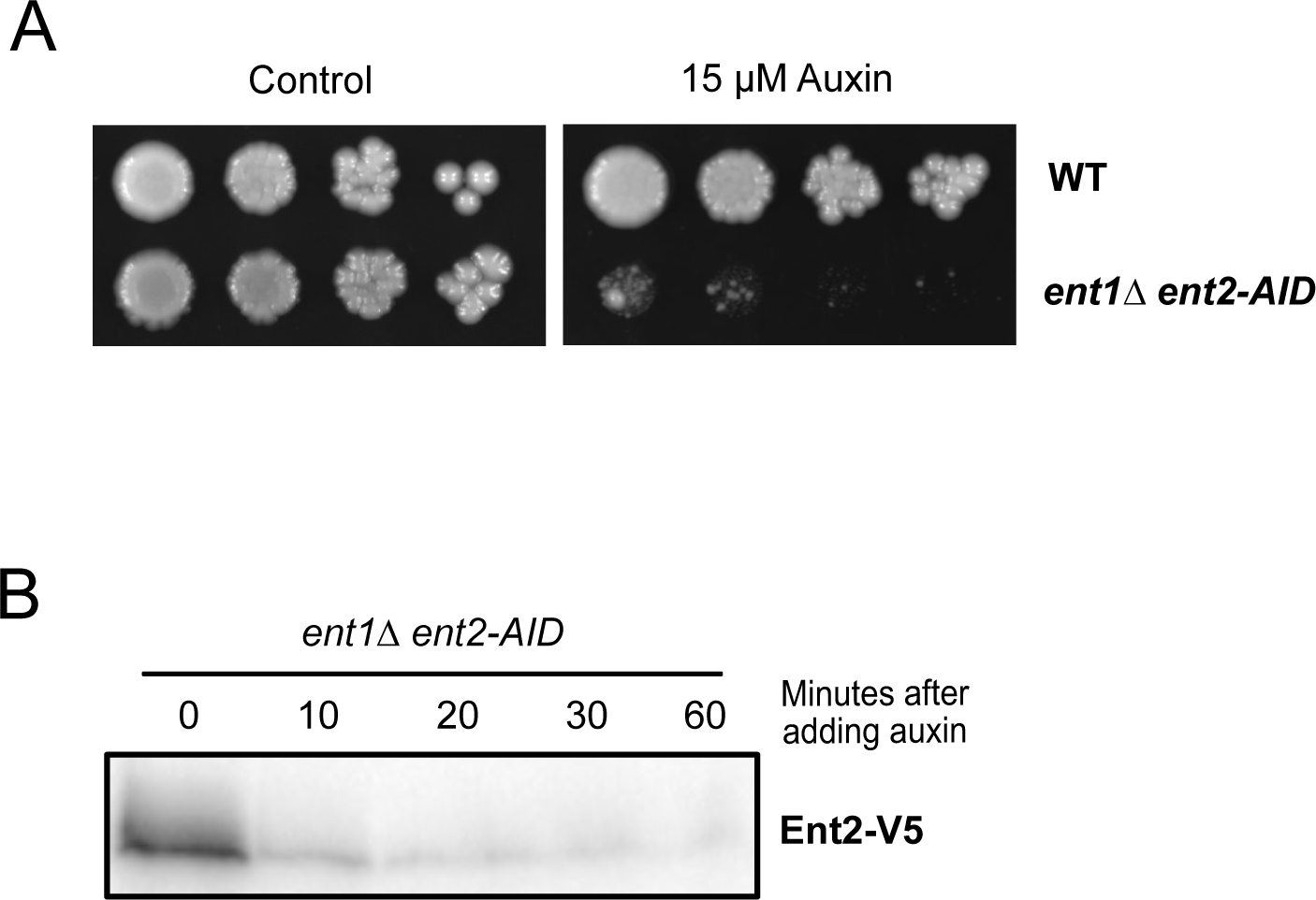
(A) A series of 10-fold dilutions of the indicated strains was grown in presence or absence of 15 µM auxin. (B) Wild type and *ent1∆ ent2*-*AID* cells were grown at 30°C to log phase in YPD medium. 1 mM auxin was added and cells were collected and processed for western blotting with an anti-V5 tag antibody to detect the ent2-AID protein at the indicated time points.

**Supplementary Figure 4.**
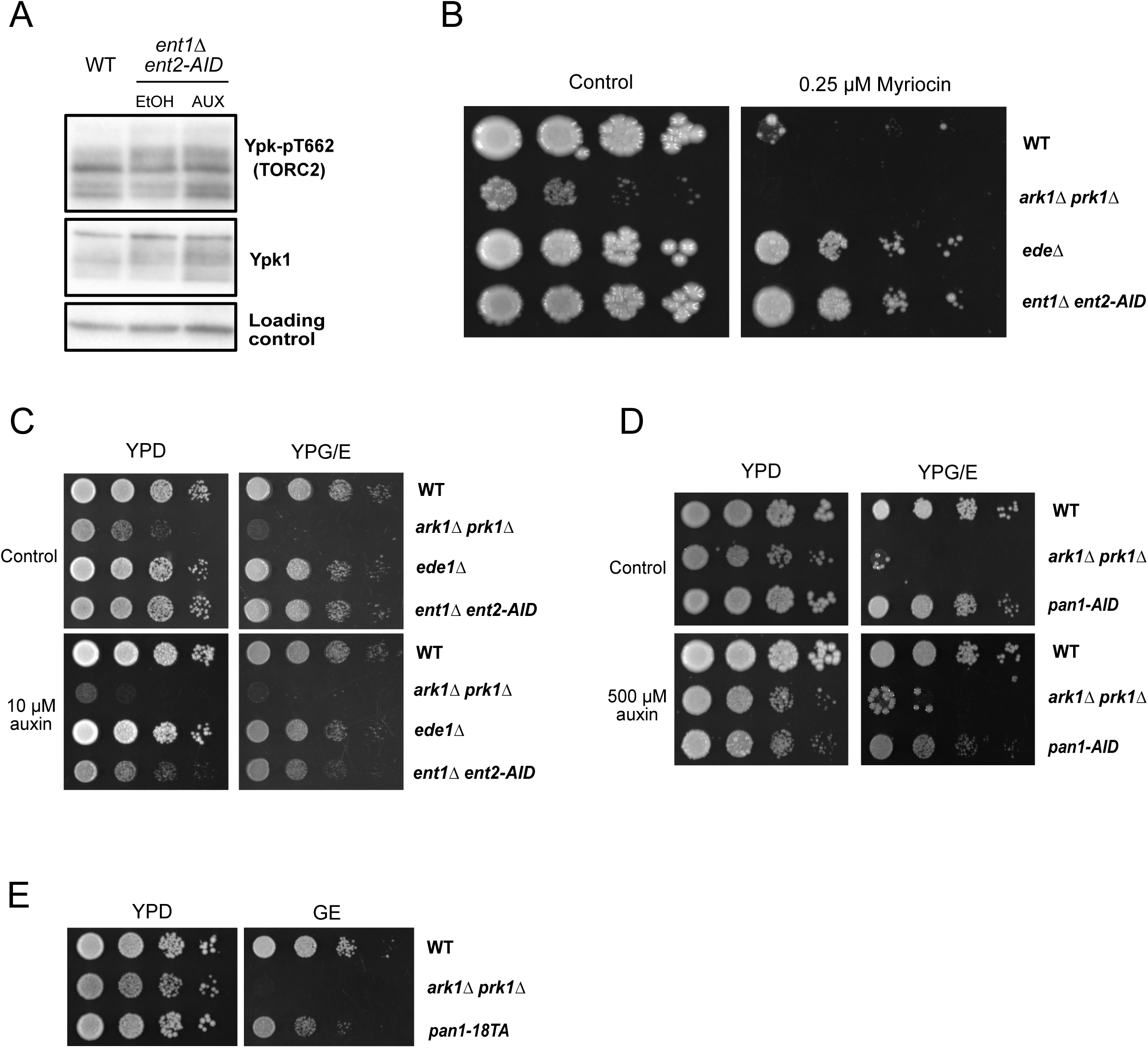
(A) Wildtype and *ent1∆ ent2*-*AID* cells were grown at 30°C to log phase in YPD medium in the presence of 10 µM auxin. As a control, the *ent1∆ ent2*-*AID* cells were also grown in the presence of the solvent for auxin (ethanol). Cells were collected and Ypk-pT662 and Ypk1 protein were analyzed by western blot. (B) A series of 10-fold dilutions of the indicated strains were grown at 30°C on YPD in the presence or absence of 0.25 µM Myriocin. (C and D) A series of 10-fold dilutions of the indicated strains were grown at 30°C on YPD or YPG/E in the absence or presence 10 µM (C) or 500 µM (D) auxin. (E) A series of 10-fold dilutions of the indicated strains were grown at 30°C on YPD or YPG/E.

## References

Alcaide-Gavilan, M., Lucena, R., Schubert, K. A., Artiles, K. L., Zapata, J., and Kellogg, D. R. (2018). Modulation of TORC2 Signaling by a Conserved Lkb1 Signaling Axis in Budding Yeast. Genetics 210, 155–170.

Artiles, K., Anastasia, S., McCusker, D., and Kellogg, D. R. (2009). The Rts1 regulatory subunit of protein phosphatase 2A is required for control of G1 cyclin transcription and nutrient modulation of cell size. PLoS Genet 5, e1000727.

Berchtold, D., Piccolis, M., Chiaruttini, N., Riezman, I., Riezman, H., Roux, A., Walther, T. C., and Loewith, R. (2012). Plasma membrane stress induces relocalization of Slm proteins and activation of TORC2 to promote sphingolipid synthesis. Nat Cell Biol 14, 542–547.

Breslow, D. K., Collins, S. R., Bodenmiller, B., Aebersold, R., Simons, K., Shevchenko, A., Ejsing, C. S., and Weissman, J. S. (2010). Orm family proteins mediate sphingolipid homeostasis. Nature 463, 1048–1053.

Casamayor, A., Torrance, P. D., Kobayashi, T., Thorner, J., and Alessi, D. R. (1999). Functional counterparts of mammalian protein kinases PDK1 and SGK in budding yeast. Curr. Biol. 9, 186–197.

Cope, M. J., Yang, S., Shang, C., and Drubin, D. G. (1999). Novel protein kinases Ark1p and Prk1p associate with and regulate the cortical actin cytoskeleton in budding yeast. The Journal of Cell Biology 144, 1203–1218.

Ferrezuelo, F., Colomina, N., Palmisano, A., Garí, E., Gallego, C., Csikász-Nagy, A., and Aldea, M. (2012). The critical size is set at a single-cell level by growth rate to attain homeostasis and adaptation. Nature Communications 3, 1012.

Gagny, B., Wiederkehr, A., Dumoulin, P., Winsor, B., Riezman, H., and Haguenauer-Tsapis, R. (2000). A novel EH domain protein of Saccharomyces cerevisiae, Ede1p, involved in endocytosis. Journal of Cell Science 113 (Pt 18), 3309–3319.

Harvey, S. L., Enciso, G., Dephoure, N., Gygi, S. P., Gunawardena, J., and Kellogg, D. R. (2011). A phosphatase threshold sets the level of Cdk1 activity in early mitosis in budding yeast. Mol. Biol. Cell 22, 3595–3608.

Heitman, J., Movva, N. R., and Hall, M. N. (1991). Targets for cell cycle arrest by the immunosuppressant rapamycin in yeast. Science 253, 905–909.

Henry, K. R., D’Hondt, K., Chang, J. S., Nix, D. A., Cope, M. J. T. V., Chan, C. S. M., Drubin, D. G., and Lemmon, S. K. (2003). The Actin-Regulating Kinase Prk1p Negatively Regulates Scd5p, a Suppressor of Clathrin Deficiency, in Actin Organization and Endocytosis. Current Biology 13, 1564–1569.

Hirsch, J., and Han, P. W. (1969). Cellularity of rat adipose tissue: effects of growth, starvation, and obesity. The Journal of Lipid Research 10, 77–82.

Johnston, G. C., Pringle, J. R., and Hartwell, L. H. (1977). Coordination of growth with cell division in the yeast Saccharomyces cerevisiae. Exp. Cell Res. 105, 79–98.

Kamada, Y., Fujioka, Y., Suzuki, N. N., Inagaki, F., Wullschleger, S., Loewith, R., Hall, M. N., and Ohsumi, Y. (2005). Tor2 directly phosphorylates the AGC kinase Ypk2 to regulate actin polarization. Molecular and Cellular Biology 25, 7239–7248.

Leitao, R. M., and Kellogg, D. R. (2017). The duration of mitosis and daughter cell size are modulated by nutrients in budding yeast. The Journal of Cell Biology 216, 3463–3470.

Leitao, R. M., Jasani, A., Talavera, R. A., Pham, A., Okobi, Q. J., and Kellogg, D. R. (2019). A Conserved PP2A Regulatory Subunit Enforces Proportional Relationships Between Cell Size and Growth Rate. Genetics 213, 517–528.

Lucena, R., Alcaide-Gavilan, M., Anastasia, S. D., and Kellogg, D. R. (2017). Wee1 and Cdc25 are controlled by conserved PP2A-dependent mechanisms in fission yeast. Cell Cycle 16, 428–435.

Lucena, R., Alcaide-Gavilan, M., Schubert, K., He, M., Domnauer, M. G., Marquer, C., Klose, C., Surma, M. A., and Kellogg, D. R. (2018). Cell Size and Growth Rate Are Modulated by TORC2-Dependent Signals. Curr. Biol. 28, 196–210.e4.

Muir, A., Ramachandran, S., Roelants, F. M., Timmons, G., and Thorner, J. (2014). TORC2-dependent protein kinase Ypk1 phosphorylates ceramide synthase to stimulate synthesis of complex sphingolipids. eLife 3, 1–34.

Niles, B. J., Mogri, H., Hill, A., Vlahakis, A., and Powers, T. (2012). Plasma membrane recruitment and activation of the AGC kinase Ypk1 is mediated by target of rapamycin complex 2 (TORC2) and its effector proteins Slm1 and Slm2. Proc. Natl. Acad. Sci. U.S.a. 109, 1536–1541.

Rispal, D. et al. (2015). Target of Rapamycin Complex 2 Regulates Actin Polarization and Endocytosis via Multiple Pathways. Journal of Biological Chemistry 290, 14963–14978.

Roelants, F. M., Baltz, A. G., Trott, A. E., Fereres, S., and Thorner, J. (2010). A protein kinase network regulates the function of aminophospholipid flippases. Proc. Natl. Acad. Sci. U.S.a. 107, 34–39.

Roelants, F. M., Breslow, D. K., Muir, A., Weissman, J. S., and Thorner, J. (2011). Protein kinase Ypk1 phosphorylates regulatory proteins Orm1 and Orm2 to control sphingolipid homeostasis in Saccharomyces cerevisiae. Proc. Natl. Acad. Sci. U.S.a. 108, 19222–19227.

Schaechter, M., Maaloe, O., and Kjeldgaard, N. O. (1958). Dependency on medium and temperature of cell size and chemical composition during balanced grown of Salmonella typhimurium. J. Gen. Microbiol. 19, 592–606.

Schneider, C. A., Rasband, W. S., and Eliceiri, K. W. (2012). NIH Image to ImageJ: 25 years of image analysis. Nat Meth 9, 671–675.

Sekiya-Kawasaki, M. et al. (2003). Dynamic phosphoregulation of the cortical actin cytoskeleton and endocytic machinery revealed by real-time chemical genetic analysis. The Journal of Cell Biology 162, 765–772.

Sun, Y., Miao, Y., Yamane, Y., Zhang, C., Shokat, K. M., Takematsu, H., Kozutsumi, Y., and Drubin, D. G. (2012). Orm protein phosphoregulation mediates transient sphingolipid biosynthesis response to heat stress via the Pkh-Ypk and Cdc55-PP2A pathways. Mol. Biol. Cell 23, 2388–2398.

Toshima, J. Y., Furuya, E., Nagano, M., Kanno, C., Sakamoto, Y., Ebihara, M., Siekhaus, D. E., and Toshima, J. (2016). Yeast Eps15-like endocytic protein Pan1p regulates the interaction between endocytic vesicles, endosomes and the actin cytoskeleton. eLife 5, 3671.

Toshima, J., Toshima, J. Y., Martin, A. C., and Drubin, D. G. (2005). Phosphoregulation of Arp2/3-dependent actin assembly during receptor-mediated endocytosis. Nat Cell Biol 7, 246–254.

Watson, H. A., Cope, M. J., Groen, A. C., Drubin, D. G., and Wendland, B. (2001). In vivo role for actin-regulating kinases in endocytosis and yeast epsin phosphorylation. Mol. Biol. Cell 12, 3668–3679.

Wendland, B., Steece, K. E., and Emr, S. D. (1999). Yeast epsins contain an essential N-terminal ENTH domain, bind clathrin and are required for endocytosis. Embo J 18, 4383–4393.

Wullschleger, S., Loewith, R., and Hall, M. N. (2006). TOR Signaling in Growth and Metabolism. Cell 124, 471–484.

Zanolari, B., Friant, S., Funato, K., Sütterlin, C., Stevenson, B. J., and Riezman, H. (2000). Sphingoid base synthesis requirement for endocytosis in Saccharomyces cerevisiae. Embo J 19, 2824–2833.

Zapata, J., Dephoure, N., Macdonough, T., Yu, Y., Parnell, E. J., Mooring, M., Gygi, S. P., Stillman, D. J., and Kellogg, D. R. (2014). PP2ARts1 is a master regulator of pathways that control cell size. The Journal of Cell Biology 204, 359–376.

Zeng, G., Yu, X., and Cai, M. (2001). Regulation of yeast actin cytoskeleton-regulatory complex Pan1p/Sla1p/End3p by serine/threonine kinase Prk1p. Mol. Biol. Cell 12, 3759–3772.

